# Glutamate methylation, a novel histone mark in diatoms: Mass spectrometry identification and structural characterization

**DOI:** 10.1101/2025.02.01.636050

**Authors:** Stéphane Téletchéa, Bérangère Lombard, Johann Hendrickx, Damarys Loew, Leïla Tirichine

## Abstract

Post-translational modifications of histones (PTMs) play a crucial role in regulating chromatin function. These modifications are integral to numerous biological processes, including transcription, DNA repair, replication, and chromatin remodeling. While several PTMs have been identified, enhancing our understanding of their roles in these processes, there is still much to discover given the potential for virtually any histone residue to be modified. In this study, we report the discovery of a novel PTM in the model diatom *Phaeodactylum tricornutum*, glutamate methylation identified by mass spectrometry at multiple positions on histone H4 and at position 96 on histone H2B. This modification was also detected in other model organisms, including *Drosophila melanogaster*, *Caenorhabditis elegans*, and humans, but not in *Arabidopsis*. Structural bioinformatics analyses, including molecular dynamics simulations, revealed that methylation of glutamate residues on histones induces displacement of these residues, exposing them to solvent and disrupting interactions with neighboring residues in associated histones. This disruption may interfere with histone complexes promoting histone eviction or facilitating interactions with regulatory proteins or complexes, which may compromise the overall nucleosome stability.

## Introduction

Chromatin refers to the molecular packaging of DNA inside eukaryotic cells. The fundamental repeating unit of chromatin is the nucleosome which consists of a histone octamer with two copies of each H2A, H2B, H3 and H4 wrapped with ∼147 bp DNA. Histones are small, positively charged proteins consisting of a large globular domain and a flexible amino-terminal tail. These proteins can undergo various post-translational modifications (PTMs), predominantly observed on the protruding histone tails, although modifications can also occur on the globular domain and the carboxyl terminus (Tropberger and Schneider, 2010). Some of the most extensively studied post-translational modifications of histones include acetylation, methylation, phosphorylation, and ubiquitination, while emerging PTMs such as S-nitrosylation, monoamylation, butyrylation, crotonylation, hydroxylation and lactylation are also gaining attention (Tan et al., 2011; Zhang et al., 2019). These PTMs play critical roles in regulating chromatin structure, ensuring not only the compaction of DNA within the nucleus, but also dynamically altering the genome’s architecture. This ongoing modulation governs the accessibility of DNA to cellular machinery, responding to a multitude of signals. PTMs are involved in a diverse array of biological processes, including transcriptional activation and silencing, cell cycle regulation, DNA repair, cell signaling, cellular differentiation, and disease regulation (Kouzarides, 2007).

Post-translational modifications of histones regulate gene expression either directly by altering the charge of histones and thus chromatin accessibility, or indirectly by recruiting enzymes and protein complexes that facilitate downstream events, often involving reader proteins (Bartke et al., 2010; Vermeulen et al., 2010). In addition to enzymes known as ‘writers’ that add modifications to histones and ‘erasers’ that remove them, ‘readers’ are effector proteins with specialized domains, such as the Tudor domain, chromodomain, PWWP domain, plant homeodomain, WD40 domain, and bromodomain (BRD), which specifically bind to modified histone residues (Hyun et al., 2017).

Our understanding of epigenetic mediated regulation of various biological processes has significantly improved with the discovery of new PTMs enabled by advancements in technologies such as mass spectrometry based proteomics. In particular, “bottom-up” nano liquid chromatography coupled to an LTQ-Orbitrap mass spectrometery (nano-LC-MS/MS) (Freitas et al., 2004; Minshull, 2014) allows for the unbiased identification of histone PTMs without the need for specific antibodies. This technology analyzes histone peptides, generated by cleaving histones into specific sequences, by examining their charge and mass for the presence of covalently attached PTMs. While many of the new post-translational modifications of histones have been identified in model organisms that have been studied for decades, newer model organisms such as *Phaeodactylum tricornutum* present opportunities to uncover novel modifications, as much remains to be discovered in these species.

While *P. tricornutum* is recognized as a well-established model organism, epigenetic research in this significant group of eukaryotic species lags behind that of other model organisms. Yet, diatoms are one of the primary groups within chromalveolates and represent some of the most abundant, diverse (including both pennate and centric taxa), and ecologically significant algae in freshwater and marine environments. They are estimated to contribute around 40% of primary production in marine ecosystems (Field et al., 1998). Diatoms are also an important source of innovation for the nanotechnology, aquaculture and pharmaceutical industries.

The comprehensive sequencing and annotation of the *Phaeodactylum tricornutum* genome (Bowler et al., 2008; Rastogi et al., 2018) revealed an atypical genetic composition resulting from successive endosymbioses and horizontal gene transfers from bacteria. The combination of genes from diverse origins has attributed them with novel and distinctive metabolic capabilities for photosynthetic organisms, including fatty acid oxidation pathways and a mitochondria-centered urea cycle (Allen et al., 2011). Previous studies have already identified a high conservation of the epigenetic machinery in *P. tricornutum* including DNA methylation and a broad array of post-translational modifications of histones (Hoguin et al., 2023; Tirichine et al., 2017; Veluchamy et al., 2013; Veluchamy et al., 2015; Wu et al., 2023; Wu and Tirichine, 2023; Zhao, 2020). This places *P. tricornutum* as an ideal model for investigating epigenetics in unicellular photosynthetic organisms, especially within an evolutionary context given its position in the tree of life and its ancestral emergence predating that of plants and animals.

In this study, we report the discovery of novel post-translational modifications on histones within the model diatom *P. tricornutum* using a nano LC-MS/MS approach. These modifications, identified in histones H4 and H2B, involve methylation of glutamate (E) at positions 52/53, 63 and 74 of histone H4 and 96 of histone H2B. We investigated the presence of these modifications in another model diatom, as well as in widely used model organisms/cells, including HeLa cells, *Drosophila melanogaster*, *Caenorhabditis elegans* and *Arabidopsis thaliana*. Due to the unavailability of functional antibodies targeting glutamate methylation, we used structural bioinformatics methods including molecular dynamics simulations to explore the effects of these modifications on histone structure within close proximity, as well as their broader impacts on nucleosome structure and stability.

## Materials and methods

### Growth conditions and histone extraction

*Phaeodactylum tricornutum* (CCMP2561, Pt1 8.6) and *Thalassiosira pseudonana* (CCMP1335) cells were cultured in artificial seawater (EASW) (Vartanian et al., 2009). The cultures were maintained at 19°C under a 12:12 light dark cycle with a light intensity of 70 µE. Cells were harvested during the exponential growth phase, at a concentration of approximately one million cells/mL. Histones were then extracted as described previously (Tirichine et al., 2014). Additionally, histones were extracted from Drosophila embryos, HeLa cells, *Arabidopsis thaliana* and *Caenorhabditis elegans* as described previously (Jufvas et al., 2011; Shechter et al., 2007).

### Protein in-gel digestion

Histone proteins were separated on 15% SDS–PAGE gels and stained with colloidal Coomassie blue (LabSafe Gel Blue™, AGRO-BIO) reagent that does not contain methanol or acetic acid. Histone bands were excised and washed, and proteins were reduced with 10 mM DTT prior to alkylation with 55 mM chloroacetamide. After washing and shrinking of the gel pieces with 100% acetonitrile (ACN), in-gel digestion was performed. All digestion were performed overnight in 25 mM ammonium bicarbonate at 30**°**C, by adding 10-20 µl trypsin at a concentration of 12.5 ng/µl (Promega). The extraction was dried in a vacuum concentrator at room temperature and redissolved in solvent A (2% ACN, 0.1% formic acid). Peptides were then subjected to liquid chromatography mass spectrometry (LC-MS/MS) analysis.

### Mass spectrometry and data analysis

Samples were analyzed by nano LC using an Ultimate3000 system (Dionex S.A.) coupled to an LTQ-Orbitrap XL mass spectrometer (MS; Thermo Fisher Scientific, Bremen, Germany). Peptides were first trapped onto a C18 column (300 µm inner diameter x 5 mm; Dionex) at 20 µl/min in 2% ACN, 0.1% trifluoroacetic acid (TFA). After 3 min of desalting and concentration, the column was switched online to the analytical C18 column (75 µm inner diameter x 50 cm; C18 PepMap^TM^, Dionex) equilibrated in 100% solvent A. Bound peptides were eluted and separated using a 0 to 30% gradient of solvent B (80% ACN, 0.085% formic acid) during 157 min, then 30 to 50% gradient of solvent B during 20 min at a 150 nl min flow rate (40°C). Data-dependent acquisition was performed in the positive ion mode. Survey MS scans were acquired in the Orbitrap on the 400-1200 m/z range with the resolution set to a value of 100 000. Each scan was recalibrated in real time by co-injecting an internal standard from ambient air into the C-trap (‘lock mass option’). The five most intense ions per survey scan were selected for CID fragmentation and the resulting fragments were analyzed in the linear trap (LTQ). Target ions already selected for MS/MS were dynamically excluded for 20 s.

Data were acquired using the Xcalibur software (version 2.0.7) and the resulting spectra where then analyzed via the Mascot^TM^ Software created with Proteome Discoverer (version 1.4, Thermo Scientific) using the in-house database containing the sequence of histone proteins from each species Supplementary Table 1. Carbamidomethylation of cysteine, oxidation of methionine, acetylation of lysine and protein N-terminal, methylation, dimethylation of lysine, arginine and trimethylation of lysine, methylation of aspartic and glutamic acid, di-glycine of lysine, phosphorylated histidine, serine, threonine and tyrosine were set as variable modifications for Mascot searches. Specificity of trypsin digestion was set and five missed cleavage site were allowed. The mass tolerances in MS and MS/MS were set to 5 ppm and 0.5 Da, respectively. The resulting Mascot files were further processed using myProMS (Poullet et al., 2007). Modified spectra were analysed manually. We only considered a spectrum valid if we clearly identified a sequence of B-ions or Y-ions and specific fragments identifying modified amino acid. Synthetic peptides (ThermoFisher Scientific) were measured with the same MS/MS method and device (LTQ-Orbitrap XL MS, Thermo Fisher Scientific) and their MS/MS spectra were compared with those of the histone peptides. The experimental protocol was designed in such a way that methanol was excluded from each step of the experimental procedure, including, histone extraction, gel staining/destaining, in-gel digestion, and LC separation. Such a procedure avoids in vitro D/E-methylation (Haebel et al., 1998). The putative E-methylated peptides were further confirmed by using extracted ion chromatogram of spiked samples with synthetic peptides and their MS/MS spectra, the gold standard for confirming a chemical identity. As D-methylated residues have a molecular weight equivalent to glutamate; therefore, either the MS/MS of a D-methylated peptide or a D-to-E-mutation peptide can explain the same MS/MS spectrum. The residue with known D-to-E-mutation was further confirmed by using synthetic peptides which contain at the same position: methylated aspartic acid (STDmeLLIR) or glutamic acid (STELLIR). No other D-methylated peptide or D-to-E-mutation peptide were identified.

### Molecular modelling

The local impact of glutamate methylation was investigated using the backrub protocol from the Rosetta 2020.37 macromolecular modelling suite (Smith and Kortemme, 2011). The 5-methyl L-glutamate structure was obtained from the Cambridge Crystallographic Data Centre (Groom et al., 2016) where the coordinates were referenced as GAVRAX after the work of Wu and collaborators (Wu, 2005). Each glutamate methylation was studied in Rosetta by generating 20 models. The full length structure of the nucleosome was taken from *Xaenopus laevis* crystallographic structure (PDB Code: 1KX5) (Davey et al., 2002) allowing to assess the structural variations and propagations of the methylation on protein side chains and backbones.

Molecular dynamics simulations of each system were performed for one microsecond using AMBER 18 (D.A. Case, 2018). The structure of *X. laevis* was taken as the reference system (PDB Code: 1KX5) (Davey et al., 2002). Three replicates of 1 microsecond were performed for the native nucleosome and 1 microsecond of simulations were computed per glutamate modification. The 5-methyl L-glutamate grafted on the reference structure was parametrized using MRP.py (Sahrmann et al., 2020). This methods allows to prepare non-standard residues by providing input files for Gaussian 16 (M. J. Frisch, 2016) and then antechamber using GAFF2 atom types. Nucleosome proteins were typed using the ff14SB force field, the DNA was typed using parmbsc1 (Ivani et al., 2016) and the water model was TIP3P (Price and Brooks, 2004). Trajectory analysis and visualization were performed using VMD (Humphrey et al., 1996), PyMOL (Schrödinger) (https://pymol.org/sites/default/files/pymol_0.xml), MDAnalysis and its HELANAL module (Bansal et al., 2000; Michaud-Agrawal et al., 2011) and matplotlib (Hunter, 2007).

## Results

### Identification of glutamate methylation in *P. tricornutum* histones

Nano LC-MS/MS analysis of *P. tricornutum* histones revealed novel post-translational modifications. Specifically, glutamate methylation was identified at residues 52, 53 (ISGLIY(EE)meTR), 63 (VFLEmeNVIR), and 74 (DSVTYTEmeHAR) of histone H4, and at residue 96 (LMoxLPGE-meLAK) of histone H2B. To validate the presence of glutamate methylation, E-methylated peptides were further confirmed using extracted ion chromatograms of spiked samples with synthetic peptides and their MS/MS spectra, considered the gold standard for chemical identification (Fig. 1). The synthetic and in vivo peptides have identical fragmentation patterns, which indicates that the methylated site is determined with high accuracy. All the positive identifications were manually inspected to ensure the quality of analysis.

**Fig. 1.**
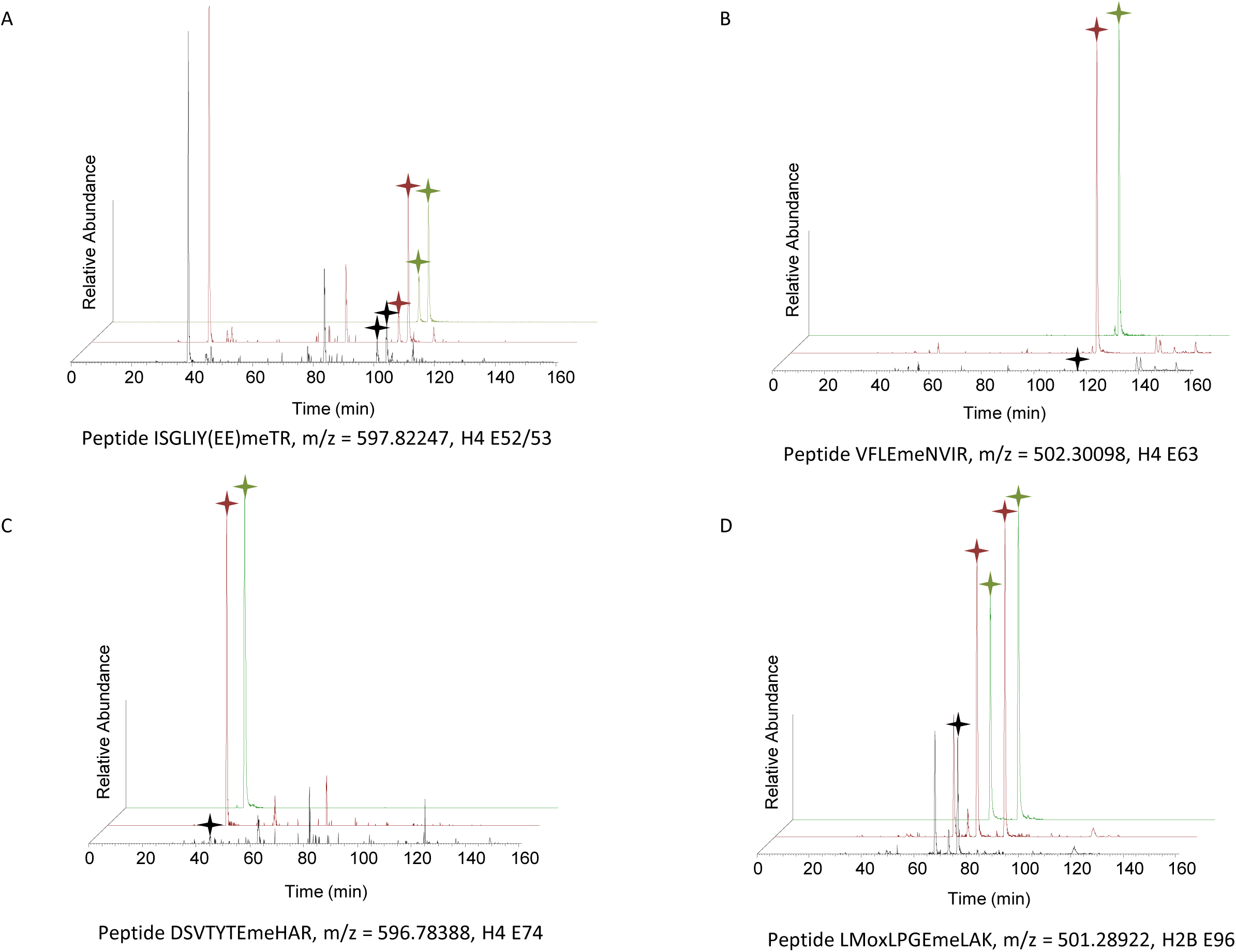
Identification and verification of E-methylated peptides in *P. tricornutum* Extracted ion chromatograms of in vivo E-methylated (black curve) histone H4 peptides at positions (A) E52 and E53, (B) E63, (C) E74, and (D) H2B peptide at position E96, compared with spiked synthetic peptides (red curve) and synthetic peptides (green curve). Stars indicate peaks identified in the tandem mass spectrum.

Glutamate methylation was observed on the globular domain of both histones (Fig. 2A), prompting questions about the implications of these modifications for nucleosomal structure and stability. This modification is localized to the histone fold domain of both H4 and H2B within the extended central α-helix, which forms homo-or heterodimers with other histone fold proteins. Methylation on both E52 and 53 as well as E63 and E74 of histone H4 are on the most solvent exposed sides of the core nucleosome while methylation on E96 of histone H2B is slightly hidden suggesting reduced interactions with binding proteins (Fig. 2B,C).

**Fig. 2.**
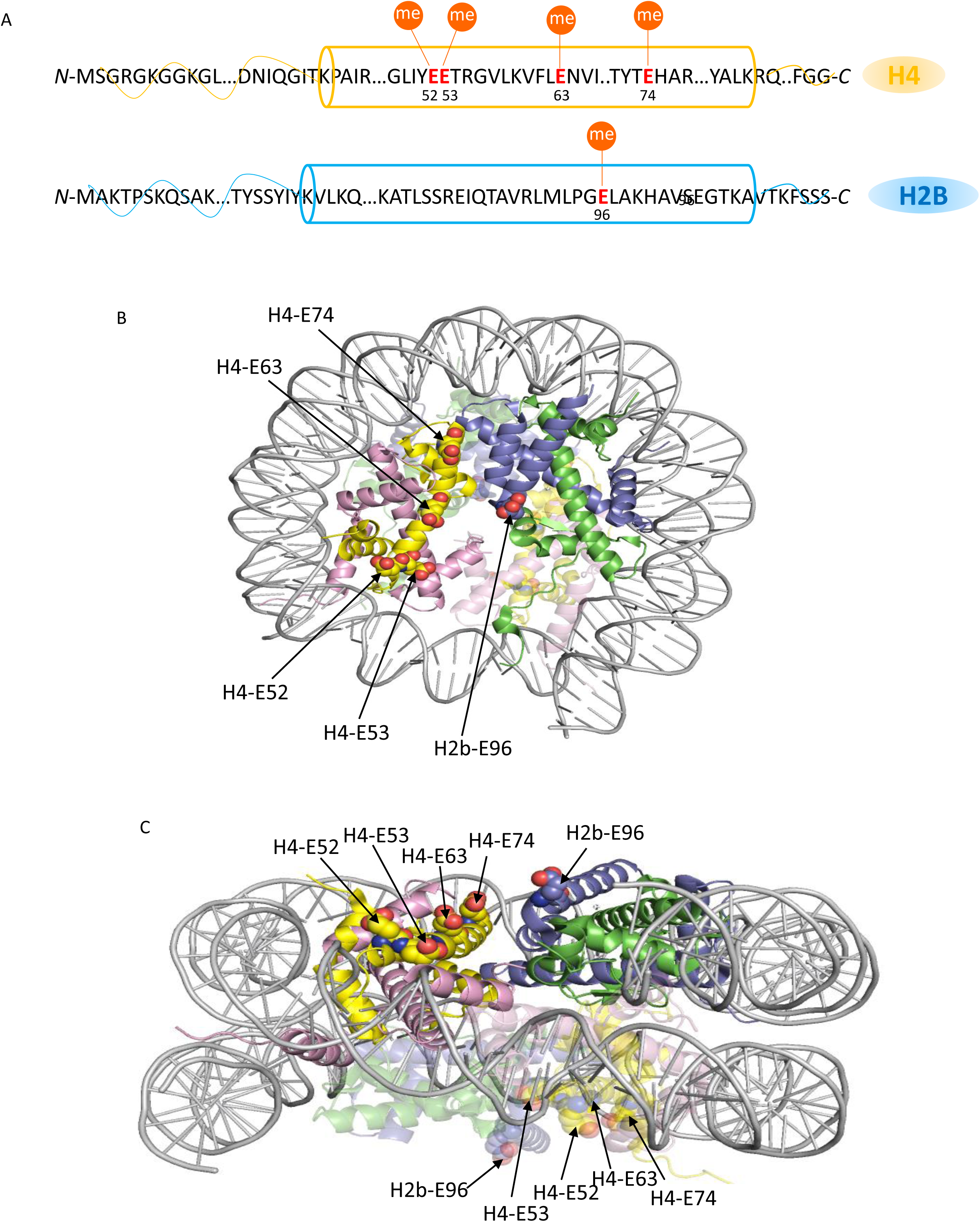
Representation of glutamate methylation on histones sequences and nucleosome. (A) Localization of glutamate methylation sites on histones H4 and H2B. The cylinder represents the histone globular domain where the modified E residues are localised. Modified residue numbers are shown below the red-highlighted E residues. Filled orange circles indicate methylation. Glutamate methylated sites are modeled on the crystal structure of the nucleosome (Protein Data Bank file 3A6N). Front (B) and side view (C) of the localization of glutamate methylation on the nucleosome. The histone proteins are shown in ribbon diagram with histone H2A in green, H2B in blue, H3 in pink, and H4 in yellow. The DNA helix is shown in gray. Modified residues are visible as red spheres. The image was generated using the program Pymol.

### Glutamate methylation on histones is prevalent across various organisms

To explore the ubiquity of E methylation across different lineages, we extracted histones from model organisms representing various branches of the tree of life. These organisms included human with HeLa cells sample, *Caenorhabditis elegans*, *Drosophila melanogaster*, *Arabidopsis thaliana col0*, and the centric diatom *Thalassiosira pseudonana*. Mass spectrometry analysis of histones from five species identified the presence of E methylation on histone H4 in *T. pseudonana* at positions E52/53 (ISGLIYEEmeTR) (Fig. 3A), and E63 (VFLEmeNVIR) of histone H4, and E96 (LPGEmeLAK) of histone H2B in *D. melanogaster* (Fig. 3B-D). In *C. elegans*, E methylation was identified at E52/53 (ISGLIY(EE)meTR) and E63 (VFLEmeNVIR) of histone H4, and E67 (AMSIMNSFVNDVFEmeR) of histone H2B. In humans, E methylation was identified at E52/53 (ISGLIY(EE)meTR), E63 (VFLEmeNVIR), and E74 (DAVTYTEmeHAK) of histone H4, and E105 (LLLPGEmeLAK) of histone H2B (Fig. 4). E methylation was not detected in *A. thaliana*, suggesting a specific function of E methylation that diatoms share with the animal kingdom but not with plants although, the possibility of its occurrence in other plant species remains to be determined.

**Fig. 3.**
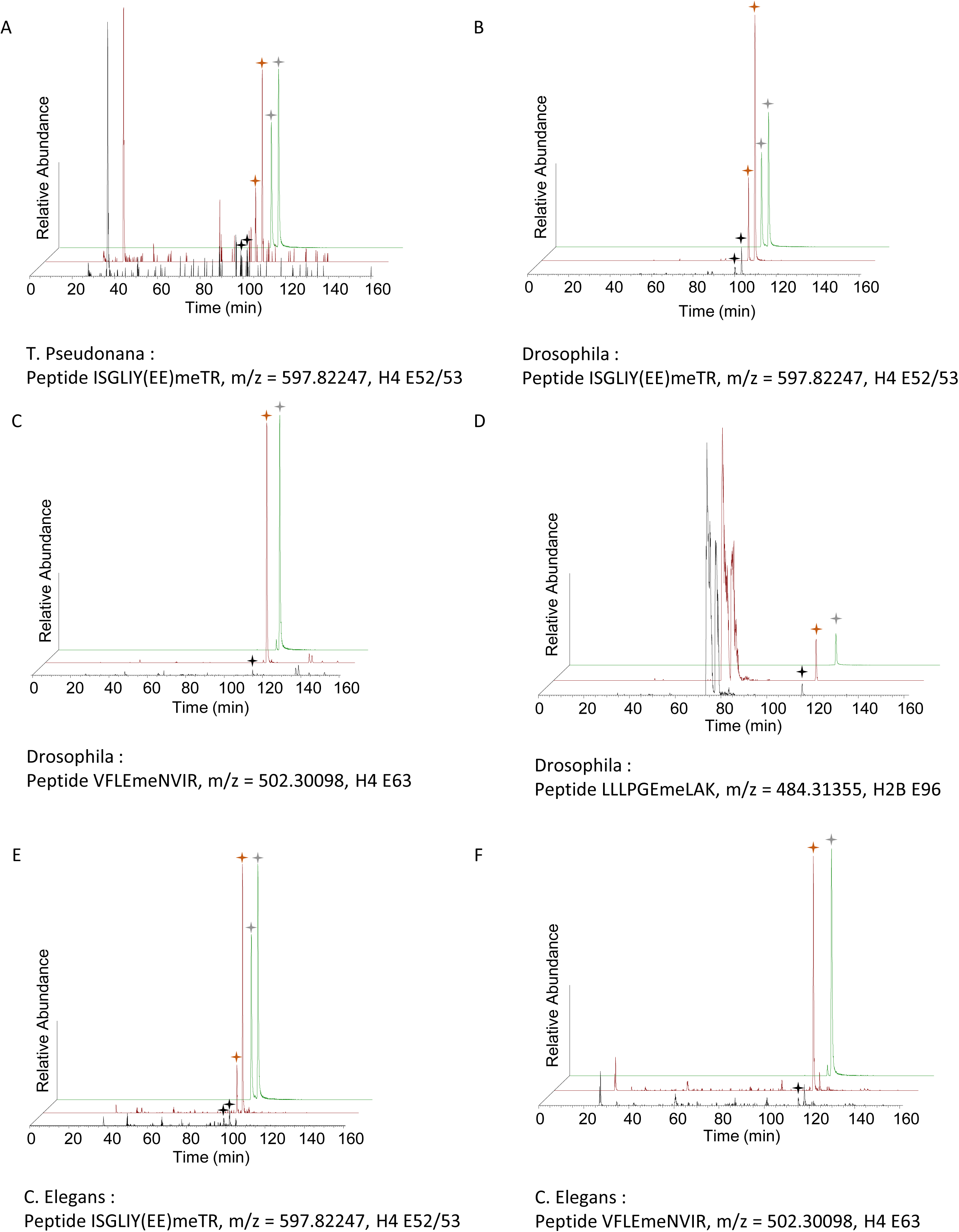
Identification and verification of E-methylated peptides in *T. pseudonana*, *D. melanogaster* and *C. elegans* Extracted ion chromatograms of in vivo E-methylated (black curve) histone H4 peptides at positions (A, B, E) E52 and E53, (C,F) E63 and H2B peptide at position (D) E96, compared with spiked synthetic peptides (red curve) and synthetic peptides (green curve). Stars indicate peaks identified in the tandem mass spectrum.

**Fig. 4.**
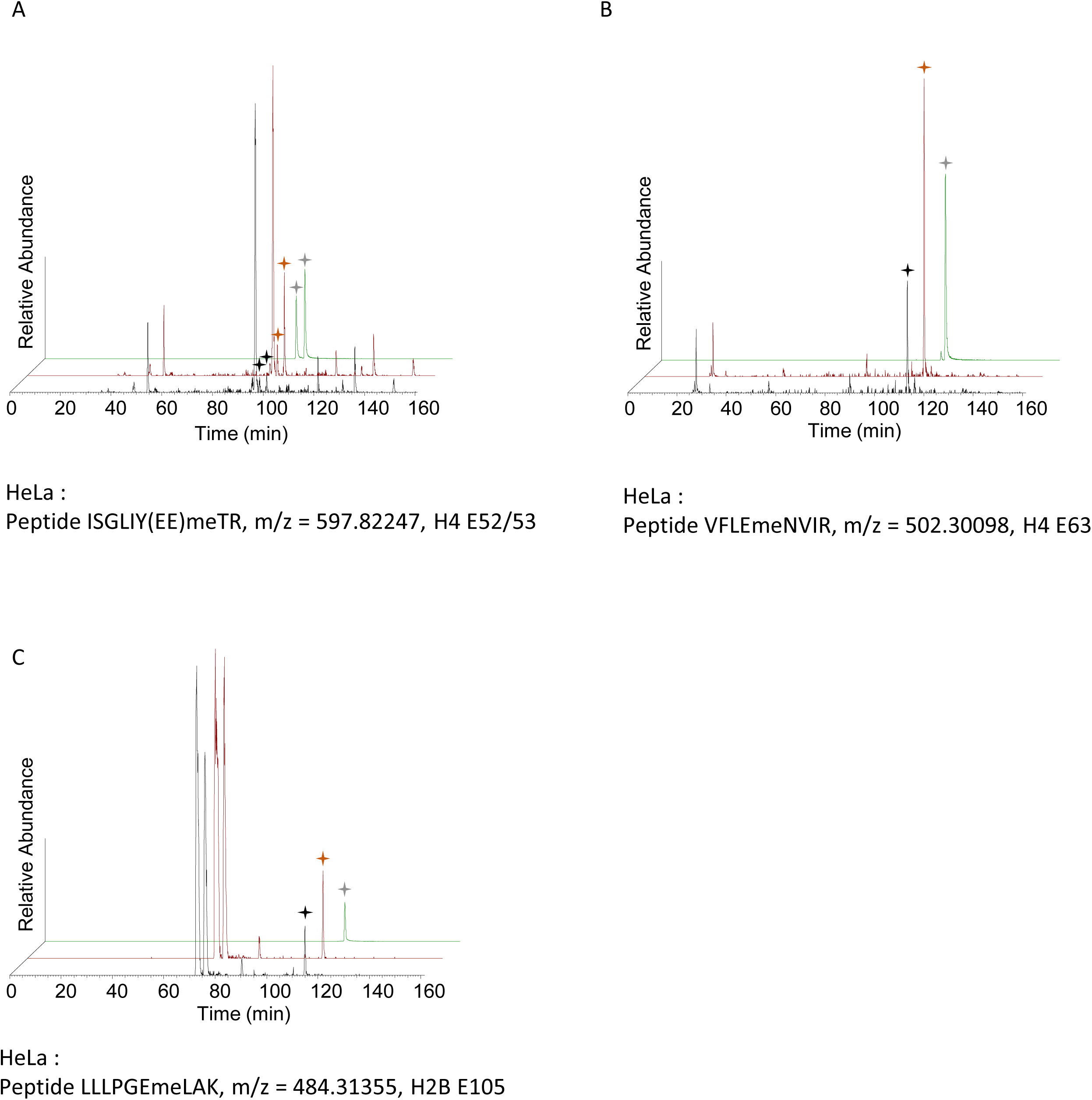
Identification and verification of E-methylated peptides in *Homo sapiens* Extracted ion chromatograms of in vivo E-methylated (black curve) histone H4 peptides at positions (A) E52 and E53, (B) E63 and H2B peptide at position (C) E105, compared with spiked synthetic peptides (red curve) and synthetic peptides (green curve). Stars indicate peaks identified in the tandem mass spectrum.

### Effects of E methylation locally, on the neighboring histones

We sought to investigate the impact of E methylation on the direct proximity of the modified histones H4 and H2B. We first studied how glutamate methylation would affect neighboring amino acids interactions using molecular modeling. In the native nucleosome, hydrogen bonds are observed between H4E52 and H4Q27, and H4E53 and H3Q125. After the methylation of E52 and E53, both bonds are disrupted, E53 keeps it position but E52 orientation changes to be more solvent-exposed (Fig. 5A). For H4E63, a salt bridge is observed within the same helix, with H4K59, and is also disrupted when E64 becomes methylated (Fig. 5B). No additional changes are observed. H4E74 is in interaction with H3K79 in the native form, but this salt bridge is disrupted when E74 is methylated, with a small rearrangement of E74 towards the solvent (Fig. 5C). H2BE96 shows two interactions, a salt bridge with K99 and a hydrogen bond with H100, two neighboring amino acids present in the next turn of their shared helix. E96 methylation leads to hydrogen bond breakage with a reorientation outwards the helix, although K99 and H100 positions seem unaffected (Fig. 5D).

**Fig. 5.**
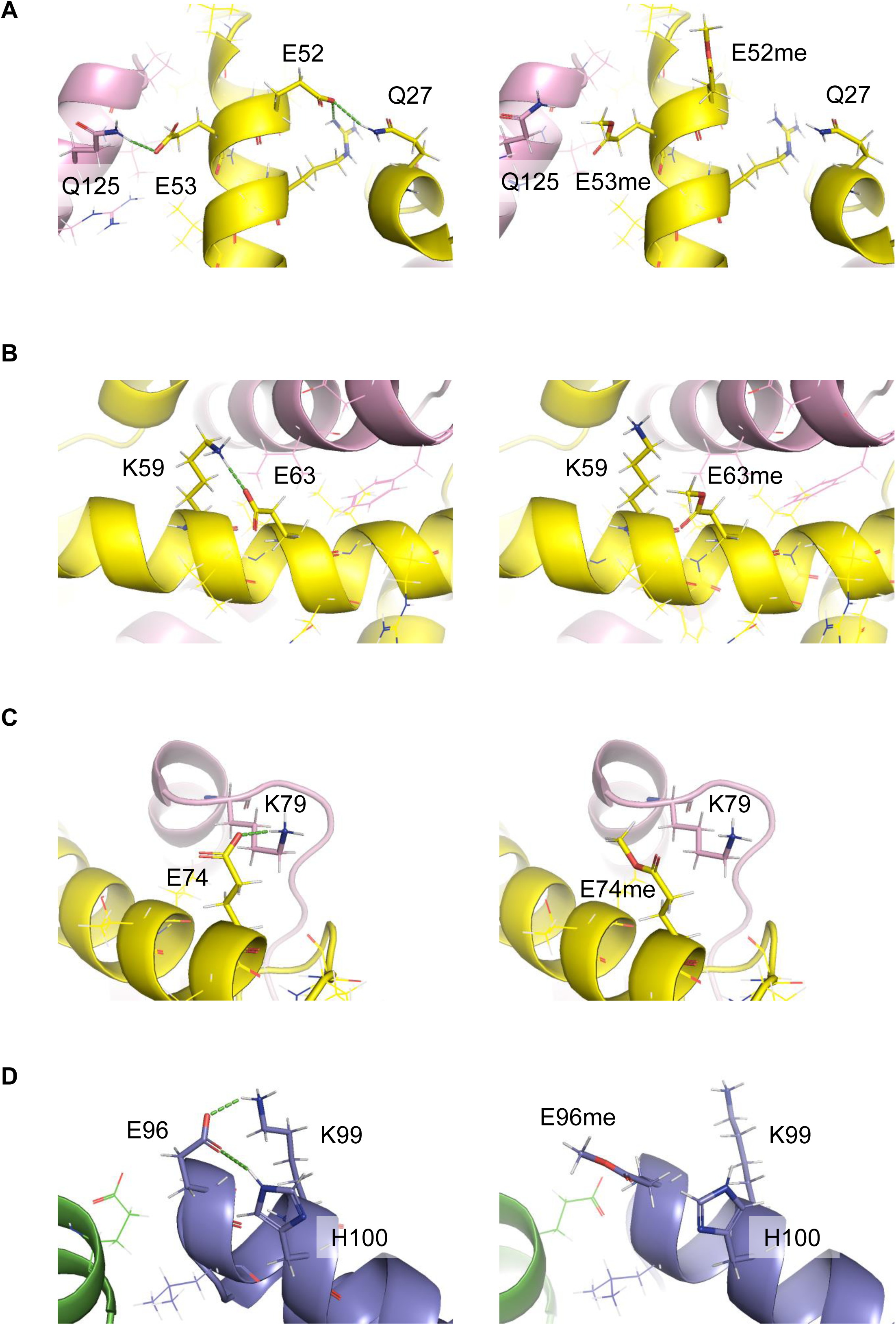
The impact of glutamate methylation at positions 52, 53, 63 and 74 of histone H4 and at position 96 of histone H2B on the adjacent amino acid residues. Comparison of amino acids interactions in the native nucleosome (left panel) or with methylated glutamate (right panel). Helices are displayed in cartoon representation, colored in yellow for H4, pink for H3, blue for H2B, and green for H2A. Hydrogen bonds between amino acids are drawn in green dashes. Hydrogens are shown as white lines, nitrogens and oxygens are shown in blue and red sticks respectively.

Out of the five positions amenable to methylation, four (E52, E63, E74, E96) showed reorientation relative to the core nucleosome structure as a result of their disrupted interactions. This movement seems however to be allowed since their location is compatible to more solvent access. In the case of H4E53, this amino acid is more buried within the nucleosome making its position more constrained by surrounding amino acids. Although, as expected, E53 lost its hydrogen bonding capacity, the methyl group addition is only correlated with a marginal shift of H3 helix position to accommodate for this bigger chemical group (Fig. 5A). This initial analysis of static structures using Rosetta allowed us to identify two distinct patterns, (i) hydrogen bond breaks with glutamate reorientation, (ii) hydrogen bond breaks without amino acid repositioning and with sterical hindrance propagating to the closer structure.

### Consequences of glutamate methylation on the histones architecture

We performed 1µs long molecular dynamics simulation to assess if observed hydrogen bond disruptions would propagate further than the backrub protocol could determine. Each modification was performed independently. To evaluate whether the observed effect also occurred in the native nucleosome, the native structure was simulated in triplicate for 1 µs. By comparing between the native and mutated states, only two amino acids display significant differences when they are methylated: (i) H4E52me, where the distance between H4E52me and H4Q27 is 9.33 +/- 2.17 Å instead of 5.83 +/- 1.56 Å, and (ii) H2BE96me where the distance between H2BE96me and H2BK99 is 7.99 +/- 1.85 instead of 5.50 +/- 2.12 in the native nucleosome (Fig. 6,7, Table S2). E96 on H2B is involved in two interactions. During the simulation time, only the interaction with K99 present a profile different from the native nucleosome simulations. No significant difference was observed with H100, the histone architecture remained intact with little deformations that could hardly be linked to local modifications.

**Fig. 6.**
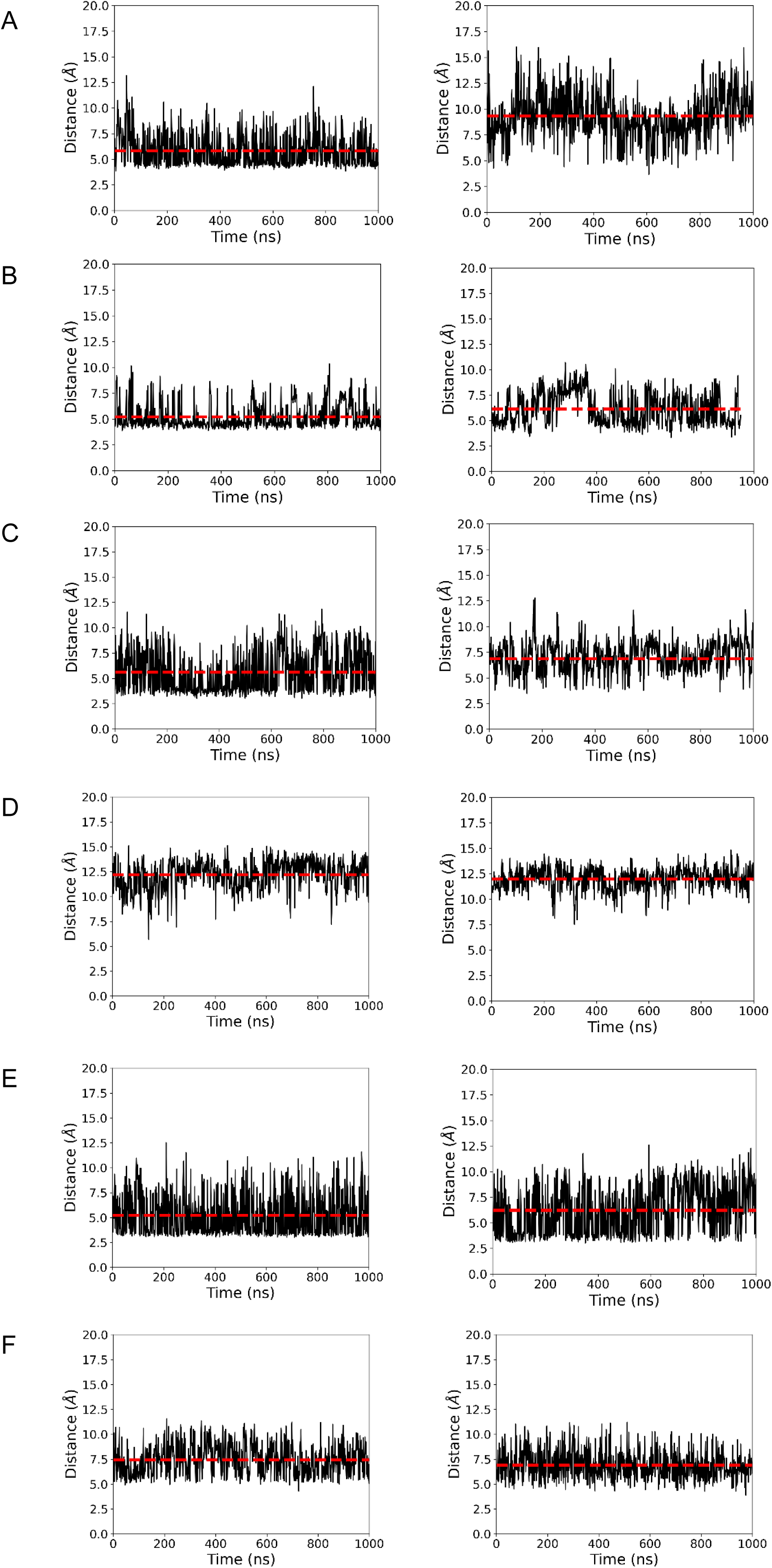
Distance over time of selected amino acids between native and methylation states. Left panels: distance observed for the native nucleosome, right panels: distance in the simulation where glutamate is methylated. (A) H4E52, (B) H4-E53, (C) H4E64, (D) H4E74, (E,F) H2B_E96.

**Fig. 7.**
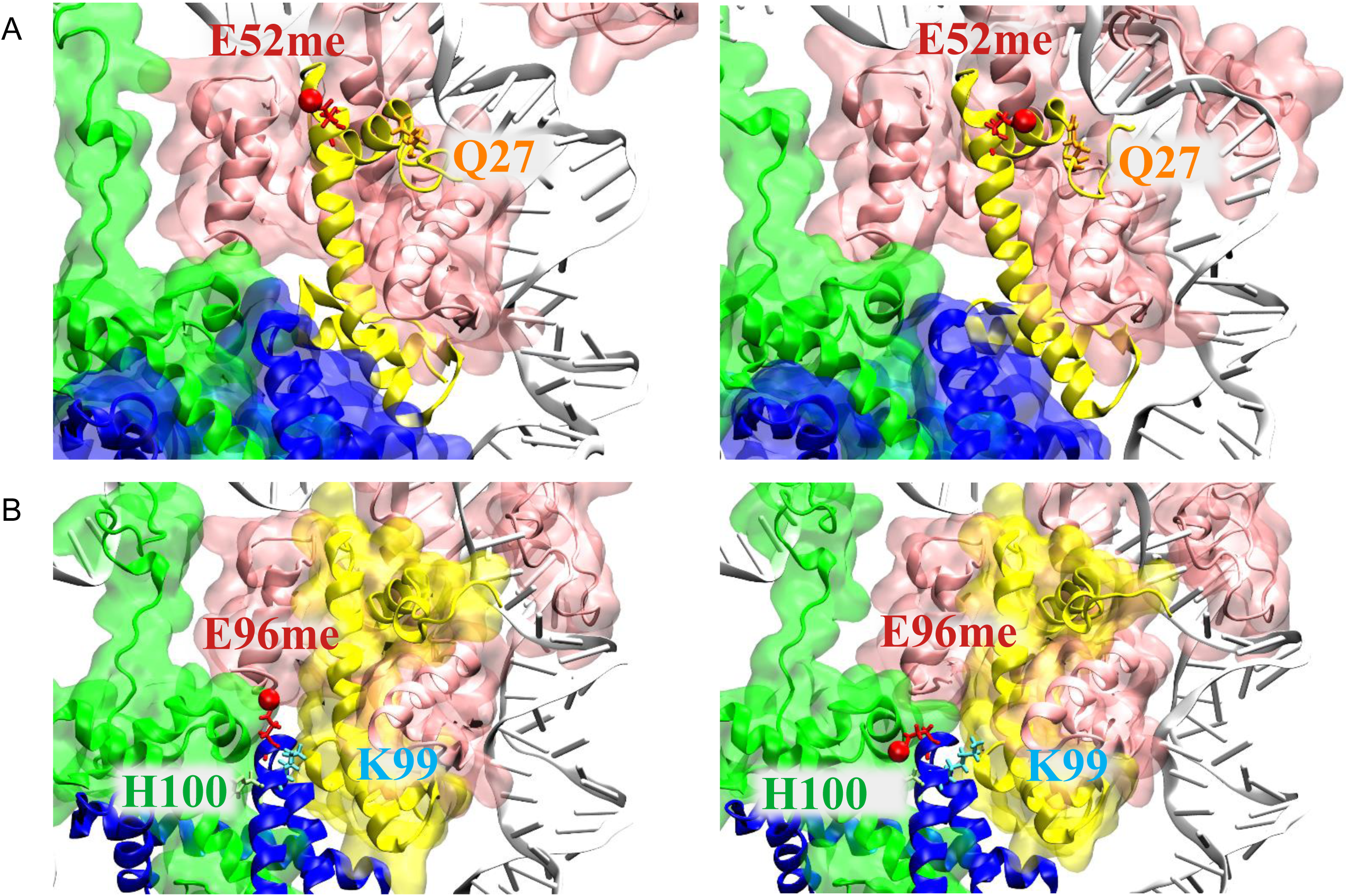
Impact of glutamate methylation on amino acids interactions. (A) H4E52 interaction with Q27. Histones are displayed in cartoon and / or surface representation, H2A in green, H2B in blue, H3 in pink and H4 in yellow, only one nucleic strand is displayed for clarity. H4E52me, the most mobile amino acid, is displayed in red sticks with the methylation position highlighted in red Van der Walls representation. (B) H2BE96 interactions with K99 (blue) and H100 (green), displayed as shown in the upper panel.

## Discussion

Mass spectrometry has proven indispensable for the direct and precise identification of histone PTMs, offering comprehensive mapping of the PTM landscape and significantly advancing our understanding of their role in regulating chromatin architecture and gene transcription. In our previous study using mass spectrometry on the model diatom *P. tricornutum*, we identified six novel positions of histone PTMs among the 65 detected (Veluchamy et al., 2015). Here, we report the identification of a novel modification, the methylation of a glutamate residue on histones H4 (positions 52, 53, 63 and 74) and H2B (position 96). These newly identified PTMs represent a significant expansion of the known repertoire of histone modifications in diatoms (Fig. 2).

To explore the extent of this histone modification beyond *P. tricornutum*, we investigated its occurrence in another model diatom, *T. pseudonana*, as well as in three other model organisms representing diverse branches of the tree of life including HeLa cells, *D. melanogaster*, *C. elegans*, and *A. thaliana* (Figs. 3,4). Our findings indicated its presence in all tested models except *Arabidopsis* suggesting a conserved function among animals but not plants although its occurrence in other plant species cannot be excluded. Given the risk of methyl esterification in vitro when methanol is used, this reagent was excluded from all experimental procedures. Furthermore, validation of glutamate methylation using synthetic peptides supports the conclusion that glutamate methylation is a bona fide modification occurring in vivo in *P. tricornutum* and all organisms tested in this study.

Glutamate methylation has been identified in mammals (HeLa cells and mice) on histones H2A (positions 67 and 95), H2B (position 35), H3 (position 59), and H4 (positions 52, 53, 63, and 74) (Zhang et al., 2018). Importantly, several positions we identified, specifically H4E52, H4E53, H4E63, and H4E74 overlap with these previously reported sites, highlighting the importance and validating the accuracy of our findings. The identification of H2BE96 is unprecedented and has not been previously reported in any organism. Furthermore, the discovery of glutamate methylation as a histone mark is novel in diatoms, flies, and worms. All the modified residues we report in this study on histones H4 and H2B in both diatom species, *C. elegans*, and *D. melanogaster* are novel PTMs that raise questions regarding their functional roles in these organisms.

Our initial analysis aimed at determining whether the local glutamate modification would globally affect nucleosome structure by identifying which interactions would be altered and whether, as a consequence, the amino acid position and/or its neighboring residues would be affected.We used an advanced backrub protocol and molecular dynamics simulations to monitor amino acids position displacements and characteristic distances and compared this measures with calculations with the native nucleosome. H4E53me was found to keep its position suggesting that despite the loss of its interaction with H3Q125, E53 remains largely in its original orientation, which may mean it is still in a relatively buried position or stabilized by other interactions in the protein. In contrast, more significant displacements were observed for the other amino acids. Specifically, H4E52me and H2BE96me induce marked reorientation of the amino acid while both H4E63me and H4E74me show more subtle changes.

Using molecular dynamics simulations, we monitored if the local deformations would propagate further in the histone or full nucleosome structure. Apart from the distances increase caused by glutamate methylation between H4E52 and H4Q27, as well as between H2BE96 and H2BK99, no other significant changes in nucleosome structure were observed. Our static and dynamic analyses indicate that the modifications result in localized rearrangements with little to no propagation, potentially facilitating nucleosome unwinding. Simulating multiple nucleosome systems both in their native state and with modifications requires substantial computational resources. Although we performed 1µs simulations, others have used simulation times an order of magnitude longer or more to capture long-range effects (Perez et al., 2012). Therefore, the length of our simulations may limit the ability to observe long-range shifts in nucleosome architecture caused by the altered orientation of methylated glutamate. A previous study extensively characterized how individual histones bind and dissociate to assemble or disrupt nucleosome structures using advanced molecular dynamics simulations, including adaptively biased molecular dynamics and umbrella sampling over much longer simulation times (Ishida and Kono, 2022). Building on this, these authors studied the energetics of distinct structural intermediates allowing them to identify key amino acids critical for DNA wrapping, unwrapping, and protein interactions. Interestingly, although our study did not establish a significant link between E74 methylation and nucleosome dynamics, Ishida et al., have identified this position as being associated with the disruption of the H2A-H2B-H3-H4 complex (Ishida and Kono, 2022). Thus, the methylations observed in our study, including E74, may be involved in core nucleosome rearrangements likely resulting in more accessible chromatin.

Furthermore, the localization of these glutamate residues within the globular domain, a region directly involved in histone DNA interactions positions them as strong candidates for interactions with other histones and DNA. It is conceivable that the increased solvent exposure of methylated E52, as a binding site, may alter the surface properties of the histone, thereby enhancing interactions with chromatin remodelers or transcription factors that promote gene expression. The hydrogen bonds between H4E52 and H3Q27, as well as between H2BE96 and H3K79, are disrupted upon methylation of E, weakening the stability of the H3-H4 octamer. This disruption may facilitate histone eviction and compromise the overall stability of the nucleosome. If we consider the charge effect, glutamate methylation decreases the negative charge of its side chain, potentially facilitating interactions with DNA. However, the distances between amino acids caused by glutamate methylation do not support close interactions with DNA, thus hindering compaction. Therefore, the effect of methylation on glutamate may stem from altered interaction dynamics rather than changes in charge.

E methylation is known to occur in bacteria modulating the chemotactic response (Shapiro and Koshland, 1993) (Clarke, 2003) as well as in eukaryotic proteins such as in yeast and HeLa cells and was reported to be abundant in eukaryotes accounting for nearly 2% of eukaryotic proteins suggesting a critical role in cellular regulation (Sprung et al., 2008). A proteomic study identified E methylation on the yeast substrate protein glyceraldehyde-3-phosphate dehydrogenase, describing it as altering three major properties of the substrate amino acid: neutralization of negative charge, increased size, and enhanced hydrophobicity. The study also compared E methylation to dephosphorylation in terms of charge change, reporting that E methylation causes more significant structural changes than the methylation of arginine and lysine, which minimally affect their side chain’s charge and hydrophobicity (Sprung et al., 2008). When identified in histones in mice, glutamate methylation levels differed between diet-induced obese mice and chow-fed controls. Specifically, there was an upregulation of H2A E67me and a downregulation of H4 E74me, suggesting a potential role in the development of obesity and diabetes.

Overall, glutamate methylation of histones appears to be a significant PTM prevalent across various organisms, playing a role in disease regulation and potentially other biological processes. The results of our study highlight the need for further investigation and comprehensive characterization of this modification. While raising antibodies against PTMs can be challenging, identifying the genomic regions targeted by glutamate methylation is crucial for understanding its function. Future research should focus on using chromatin immunoprecipitation with deep sequencing (ChIP-Seq) to identify these targets. Integrating ChIP-Seq data with transcriptomics will reveal whether this modification acts as a repressive or permissive mark. Additionally, the enzymes responsible for adding, removing, and recognizing this modification have yet to be discovered, and identifying them will contribute to our understanding of its functional mechanisms. Given the complexity of epigenetic regulation, it is also important to investigate the interactions between glutamate methylation and other epigenetic marks, such as DNA methylation and other PTMs.

## Acknowledgements

We thank the GLiCID Computing Facility (Ligerien Group for Intensive Distributed Computing, https://doi.org/10.60487/glicid, Pays de la Loire, France) for providing the bioinformatics facilities used in this work.

## Author contributions

LT conceived and designed the study. ST designed the bioinformatics analysis. LT extracted histones and ran gels for mass spectrometry. BL prepared samples for mass spectrometry and ran the mass spectrometry analysis. DL supervised the mass spectrometry analysis. ST and JH performed the bioinformatics analysis. All authors analysed, interpreted and discussed the results. LT coordinated the study and wrote the manuscript with input from all authors.

## Data availability

The data supporting the conclusions of this study are available in the paper and in the online supplementary materials. The mass spectrometry proteomics data have been deposited in the PRIDE database, a partner repository of the ProteomeXchange Consortium, with the dataset identifier PXD053679.

---

**Fig. S1.**
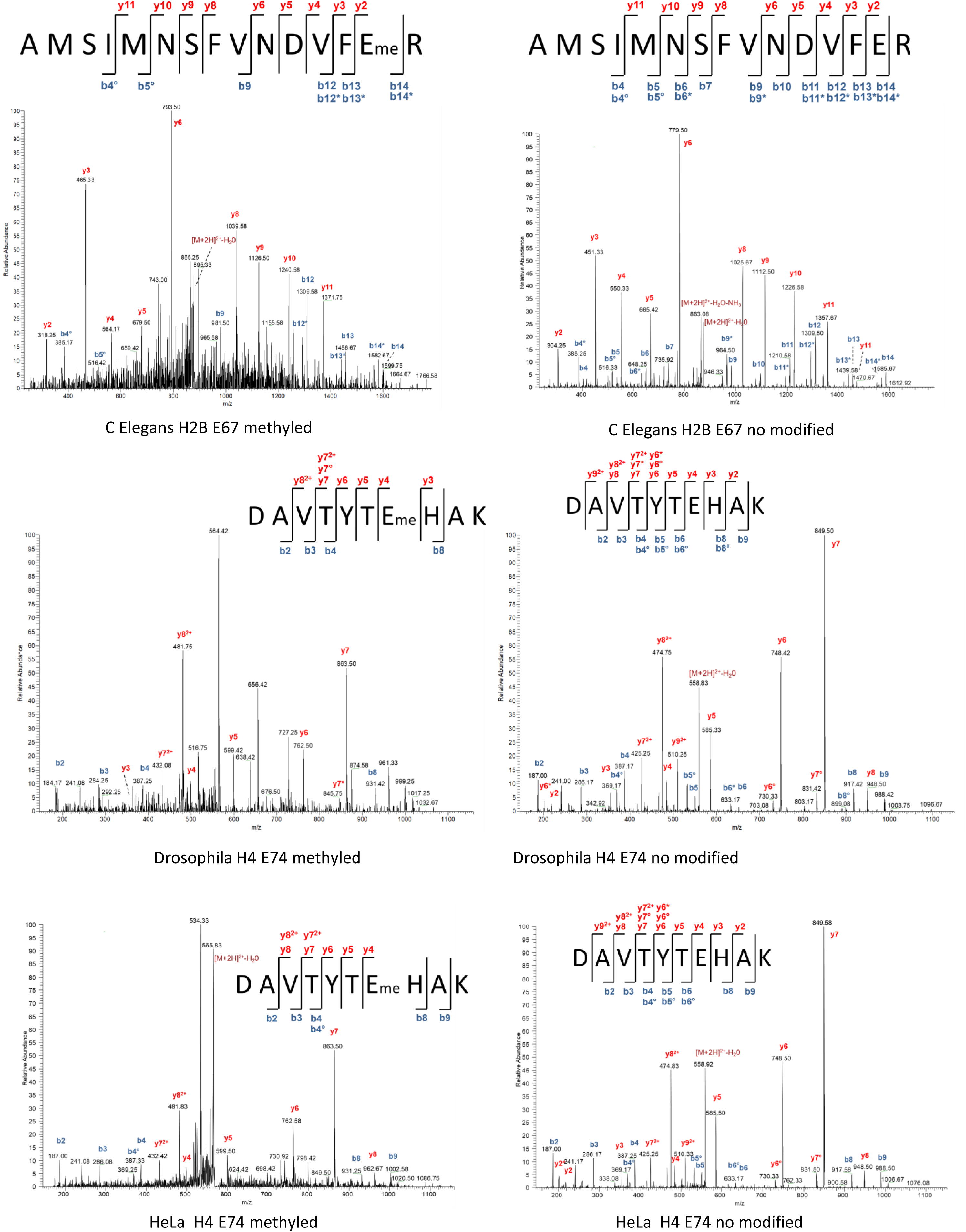
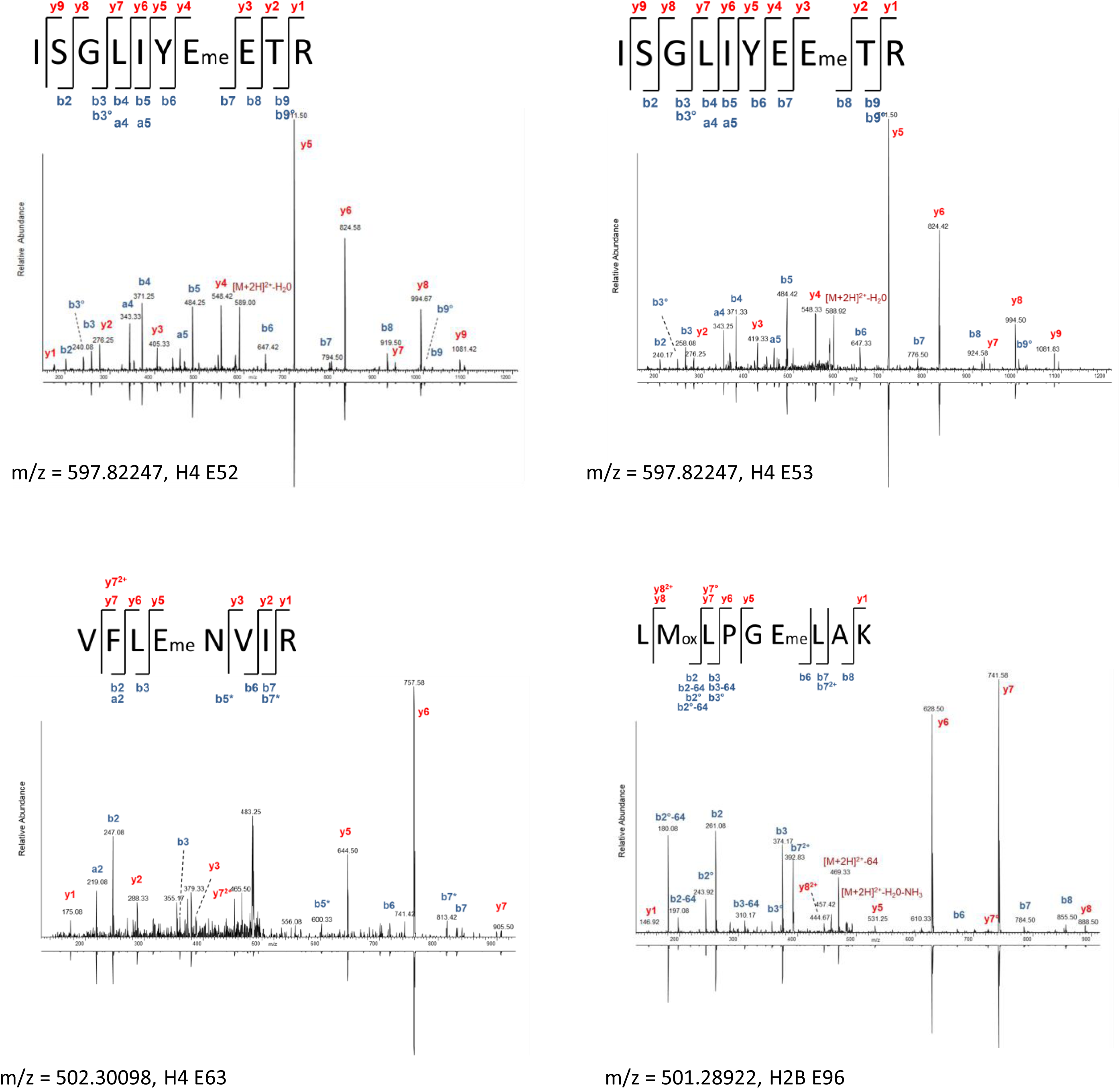
MS/MS spectra of E-methylated residues of histones H4 and H2B across different species. MS/MS spectra are derived by collision-induced dissociation of the (M+2H)^2+^ precursor. Eme indicates methylated glutamic acid.

**Table S1**. Identification number of the histones analysed in the study.

**Table S2**. Distances between heavy atoms over time for native and methylated nucleosomes.

